# Microelectrode array scaled for human hippocampal slices

**DOI:** 10.1101/2025.02.27.640534

**Authors:** Anssi Pelkonen, Vera Lezhneva, Tomi Ryynänen, Joose Kreutzer, Valeriia Sitnikova, Mireia Gómez-Budia, Adriana Della Pietra, Antonios Dougalis, Jenni Kyyriäinen, Mastaneh Torkamani-Azar, Henri Eronen, Tuomas Rauramaa, Arto Immonen, Ville Leinonen, Alejandra Sierra, Reetta Kälviäinen, Tarja Malm

## Abstract

Temporal lobe epilepsy (TLE) is a prevalent neurological disorder characterized by recurrent seizures originating from the cortex, amygdala and especially hippocampus. While two-thirds of TLE patients achieve seizure control through medication, approximately one-third remain refractory to pharmacological interventions. For these individuals, surgical resection offers a potential curative option, with approximately 70% achieving seizure freedom. However, the pathogenesis of TLE remains incompletely understood, necessitating further investigation. Therefore, resected brain tissue obtained during the surgery provides a valuable resource for *ex vivo* study of pathological neuronal activity. Currently, microelectrode array (MEA) technology is widely used for electrophysiological studies. However, commercially available MEAs are limited in their ability to record from large tissue samples, such as an entire hippocampal section. To address this limitation, we have developed a custom MEA and sample chamber compatible with commercially available headstages. This system enables recording of extracellular action potentials (EAPs) and local field potentials (LFPs) across human hippocampal tissue, providing a valuable tool for investigating the neurophysiological mechanisms underlying TLE.

**Graphical abstract:** 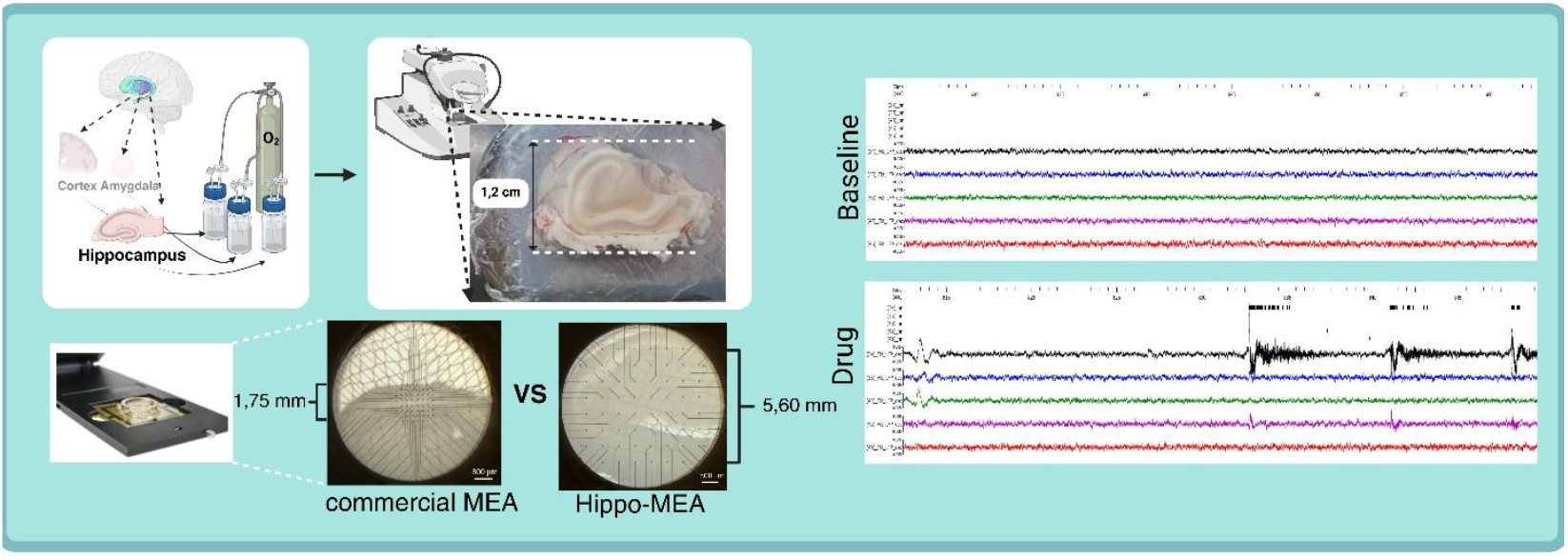

**Highlights:**

- A custom microelectrode array and chamber created for human hippocampal slices
- The custom array recorded action potentials from human hippocampal slices
- The custom array recorded local field potentials from human hippocampal slices

## 1. Introduction

Excisions from different areas of the brain from patients with epilepsy are a valuable source of living human brain samples, especially for the hippocampus. However, monitoring the network-wide electrophysiological activity of these samples is hindered by most tools being more suitable for animal models. This is especially evident with microelectrode arrays (MEAs) where the typical electrode area (2.7 mm^2^) is big enough for rodents, but not for human hippocampi (Fan et al., 2019). MEAs have been used to record human hippocampal slices in the past, but the commercially available models cover only a limited area (Hsiao et al., 2015; Wickham et al., 2020).

The hippocampus is considered as the main structure responsible for seizures initiation in epilepsy (Buckmaster et al., 2022; Jamiolkowski et al., 2024). Moreover, the most frequently observed structural pathology of temporal lobe epilepsy (TLE) is mesial temporal sclerosis which is characterized by loss of neurons in several hippocampal areas (Che et al., 2024), making this part of the brain highly interesting for epilepsy studies. The hippocampus consists of multiple subfields (CA1–CA4), the pathological interactions of which have been shown in multiple TLE studies (Núñez-Ochoa et al., 2021; Oliva et al., 2023; Reyes-Garcia et al., 2018; Whitebirch et al., 2022). Considering the complexity of hippocampus, it is highly important to record neuronal signals from multiple regions simultaneously to get the whole picture of the pathological circuits’ activity. In our ongoing consortium project, we explore the multimodal characteristics of TLE patients’ brain, where one part of the study includes MEA recordings during drug applications.

The few MEA designs with larger electrode areas (up to 42.6 mm^2^) and electrode densities have many advantages but are so customized that they cannot be combined with widely available data acquisition hardware or software (Suzuki et al., 2023; Tsai et al., 2017). They also have opaque electrode areas limiting options for microscopy or have non-optimal sample chamber designs. Here we have developed a custom-made MEA entitled Hippo-MEA where 60 electrodes are spread out to an area of 5.6 mm x 5.6 mm (31.4 mm^2^), which is sufficient to sample different subfields across the human hippocampus. The MEA is equipped with a custom-made chamber for the sample (named Sample Cup) big enough to encompass the whole slice. The benefit of the design is not only that it can encompass a large sample, but the MEA chip is made compatible with a commercially available headstage that is already widely in use in electrophysiology laboratories.

## 2. Materials and methods

### 2.1. MEA production, assessment of MEA properties and sample chamber

The MEAs (Fig. 1A-D) were built on 49 mm × 49 mm × 0.7 mm borosilicate glass slides (g-materials, Deggendorf, Germany) and had titanium tracks, titanium nitride-coated electrodes and contact pads made by the in-house ion beam-assisted -e-beam deposition (IBAD) process described in (Ryynänen et al., 2018) and silicon nitride as the passivation layer. The size of the glass plates, electrode count and contact pad placements were designed to be compatible with 60-electrode headstages from Multi Channel Systems (Reutlingen, Germany). Sixty round electrodes, θ 60 µm, were arranged in an 8×8 grid with an electrode-to-electrode pitch of 800 µm. Electrode 15 (i.e. electrode at column 1, row 5) was omitted from all analyses as it is mapped together with the reference electrode in the recording system. Electrode impedance was measured as described earlier (Ryynänen et al., 2018). Root-mean-square (RMS) noise of unfiltered electrode data (acquired at 20 kHz) was assessed from 30 s of unfiltered data acquired during normal baseline recording conditions (described below) using the following formula:

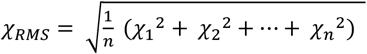

**Fig. 1.**
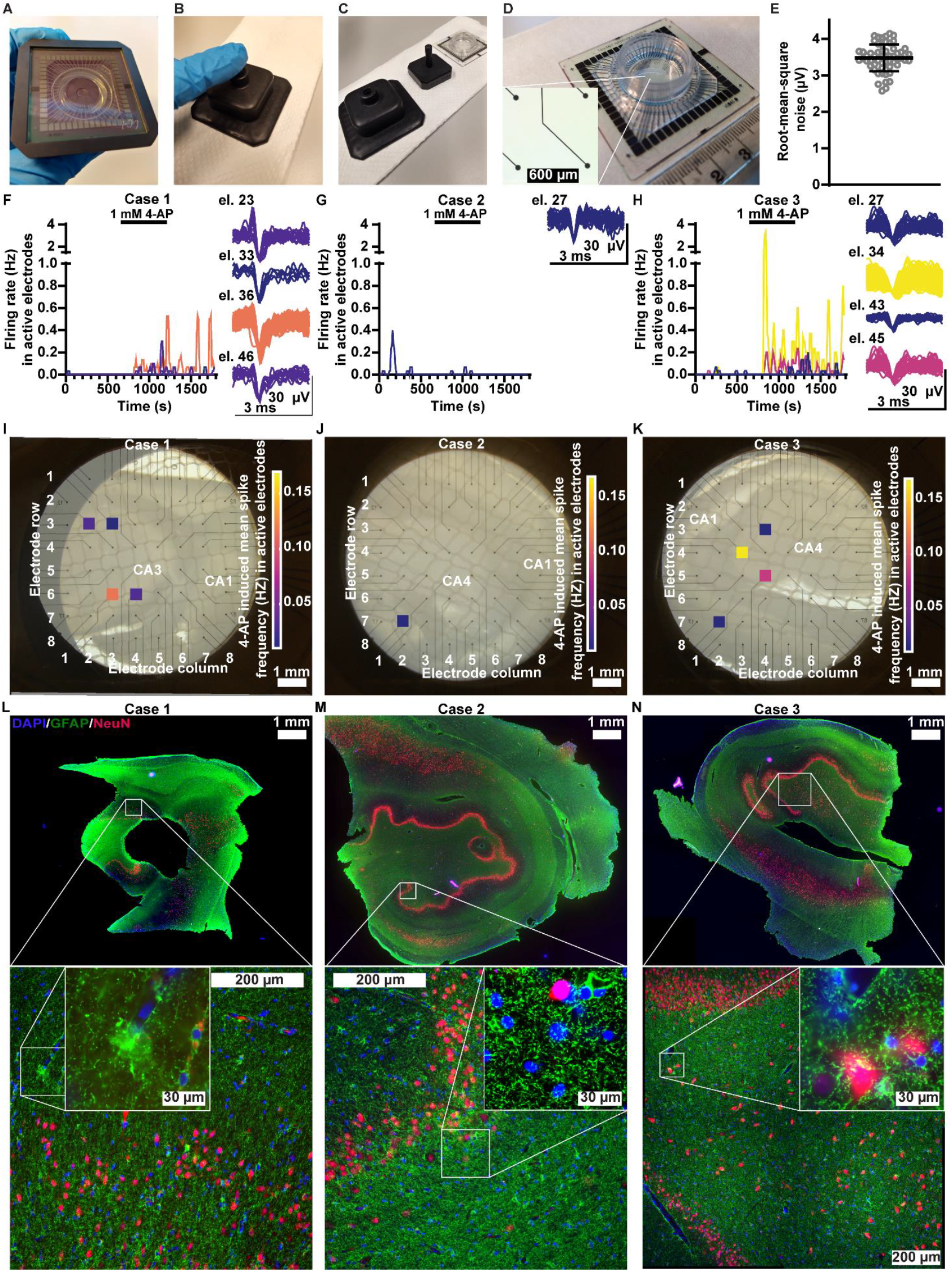
Hippo-MEA assembly, recording EAP activity and IHC. **A**, Silicone-made Sample Cup and Hippo-MEA are aligned with the Mounting Tool. **B**, Hippo-MEA and Sample Cup are bonded by pressing them gently together using the Mounting Tool. **C**, The outer and inner part of the Mounting Tool and assembled Hippo-MEA. **D**, Assembled Hippo-MEA with centimeter scale. Closeup of the electrode area in the inset (electrode pitch 800 µm). **E**, Root-mean-square noise of the 59 recording electrodes during baseline recording. **F, G, H**, EAP firing rate in active electrodes over time in 10 s bins in cases 1 (F), 2 (G) and 3 (H). Each active electrode’s data is plotted separately. Overlays of the detected EAPs in the active electrodes are shown on the right in each subfigure. In the electrode designations the first digit refers to the electrode column and the second to the row. **I, J, K**, positioning of the active electrodes in cases 1 (I), 2 (J) and 3 (K) on top of a slice-electrode overlay image. Color code indicates the average spike frequency in the electrode during maximal 4-AP effect. Slice orientation indicated by marking some of the hippocampal subfields (CA). Please note that the slice from case 1 represents rostral hippocampus, whereas the slices from cases 2 and 3 are more caudal. **L, M, N**, IHC staining for nuclear (DAPI, blue) neuronal (NeuN, red) and astrocytic (GFAP, green) markers of slices near the Hippo-MEA slice in cases 1 (L), 2 (M) and 3 (N). The stained section from case 1 is incomplete due to the original IHC slice being thinner (300 µm) than in the other cases (990 µm).

The silicone-made Sample Cup for Hippo-MEA had an inner diameter of 17 mm and wall height 9.5 mm (BioGenium Microsystems Oy, Tampere, Finland). The Sample Cups were mounted on top of the MEAs utilizing a specific device, the Mounting Tool (Fig. 1A-D). It assists with aligning and bonding the Sample Cups to the center of the MEAs, and on the other hand, reduces human errors in misaligning the Sample Cup on top of the MEA’s contact pads.

### 2.2. Patients and surgery

Hippocampal slices were obtained from patients undergoing neurosurgical tissue resection to treat TLE. The surgeries were performed as described earlier (Jutila et al., 2014, 2002). Patients were under balanced general anaesthesia induced by propofol, remifentanil, rocuronium and fentanyl. All patients had given their informed consent for the use of resected tissue prior to surgery and the study followed the Code of Ethics of the World Medical Association (declaration of Helsinki). The Ethics Committee of the Kuopio University Hospital has approved the study (128/2015, 21.4.2015). Details of patients are described in table 1.

**Table 1.** Patient information. Information is limited for information minimization purposes. ATL = anterior temporal lobe, HS = hippocampal sclerosis, FCD = focal cortical dysplasia. Internation League Against Epilepsy (ILAE) histopathological classifications: (Blümcke et al., 2013; Najm et al., 2022).

| Case | Sex | Age at surgery (years) | Type of Surgery, Side | Histopathology |
| --- | --- | --- | --- | --- |
| 1 | Male | 11–15 | ATL resection; Right | HS-ILAE1 and ATL FCD ILAE IIIa |
| 2 | Male | 31–35 | ATL resection; Left | HS-ILAE2 and ATL FCD ILAE IIIa |
| 3 | Female | 41–45 | ATL resection; Left | HS-ILAE2 |

### 2.3. Sample transport and preparation

Resected tissue samples were placed in transport bottles, each filled with 250 ml of ice-cold constantly oxygenated (95 % O_2_ / 5 % CO_2_) N-methyl-D-glucamine-based artificial cerebrospinal fluid (aCSF, exact solution compositions at (Fagerlund et al., 2022; Gazestani et al., 2023)). The samples were delivered to the laboratory within 15 min and sliced in ice-cold oxygenated aCSF. Thickness of the coronal slices was 300 µm for the Hippo-MEA recordings. The slices were kept in recovery solution (Fagerlund et al., 2022; Gazestani et al., 2023) for 45 min at +34 °C and 8-10 h at room temperature before Hippo-MEA recordings.

### 2.4. Electrophysiological recordings and analysis

The basic MEA recording protocol and recording solution is described in (Fagerlund et al., 2022; Gazestani et al., 2023). Data from the Hippo-MEA was recorded using the MEA2100-Mini-system (Multi Channel Systems, Reutlingen, Germany). An EP2 Ag/AgCl pellet (World Precision Instruments, Sarasota, FL, USA) was used as a reference electrode. Experiments were performed under constant perfusion with recording solution (3 ml/min, 32-34 °C, solution oxygenated with 95 % O_2_/5 % CO_2_ gas) and the slice was kept in place with a HSG-MEA-5AD anchor (ALA Scientific, Farmingdale, NY, USA). The slice was allowed to settle for 30 min on MEA before recordings. The actual recording consisted of a 10 min baseline, 10 min treatment with 4-aminopyridine (4-AP; 1 mM; 10122610, Thermo Scientific Chemicals, Waltham, MA, USA) and a 10 min washout.

Analysis of extracellular action potential (EAP) and local field potential (LFP) data was based on a NeuroExplorer (Plexon, Dallas, TX, USA) pipeline described earlier (Fagerlund et al., 2022). In brief, EAPs were detected from band-passed filtered signal (300-3000 Hz, 2nd order Butterworth) when their amplitude exceeded -5.5 times standard deviation (SD) of noise. Electrodes reaching the active electrode criteria (min. two EAPs, mean frequency 0.05 Hz, instantaneous frequency 0.1 Hz) in any 10 s bin during the peak 4-AP effect (970 s to 1270 s) were considered to display true EAP activity. The LFP data was extracted from low-pass filtered signal (<200 Hz, 2nd order Butterworth) for five frequency bands: delta (δ) 1–3 Hz, theta (θ) 4– 8 Hz, alpha (α) 9-13 Hz, beta (β) 14-30 Hz and gamma (γ) 30–100 Hz. A threshold-method was used to identify the electrodes responding at each frequency range of the LFP data. The power spectral density (PSD) at baseline (0 s to 600 s) was averaged for each electrode and frequency range, determining noise at baseline. If the PSD at any 10 s bin during the 4-AP period (670 s to 1270 s) exceeded 5.5 times SD of the baseline noise, the electrode was considered as responding.

### 2.5. Immunohistochemistry

To ensure intact cryosections for stainings, the coronal immunohistochemistry (IHC) slices were 990 µm thick, apart for case 1 where only 300 µm slices were available. The rostral-caudal distance of the IHC slice to the Hippo-MEA slice in tissue was 300 µm in case 1, 1500 µm in case 2 and 0 µm in case 3. The samples were fixed, cryoprotected, cryosectioned into 20 µm thick sections and stained with methods described earlier (Jäntti et al., 2022). The primary antibodies used were: NeuN (MAB377; Milipore, Darmstadt, Germany; 1:200 dilution) for neuronal visualization and GFAP (Z0334; DAKO, Glostrup, Denmark; 1:750 dilution) for astrocyte staining. Alexa Fluor 488 (A11008; Life Technologies, Carlsbad, CA, USA; 1:300 dilution) and Alexa Fluor 568 (A11004; Life Technologies; 1:300 dilution) were used as secondary antibodies. Images were obtained with Leica Thunder Imager 3D Tissue Slide Scanner (Leica Microsystems, Wetzlar, Germany).

## 3. Results and discussion

### 3.1. Hippo-MEA properties

Impedance of electrodes in Hippo-MEA was 35±5 kΩ (mean±SD), lower than reported for electrodes produced with the same method earlier (83±5 kΩ)(Ryynänen et al., 2018). While a lower impedance is generally a sign of better signal detection, the measured value here is due to a larger electrode diameter (60 µm vs. 30 µm). During baseline recording the RMS noise was 3.4±0.5 µV (Fig. 1E) which is similar to electrodes produced with the same method earlier (Ryynänen et al., 2018). The Mounting Tool enabled easy assembly of the Sample Cup and the Hippo-MEA (Fig. 1A-D). Silicone device attachment to glass-like MEA surfaces can require e.g. plasma treatment (Pelkonen et al., 2020) but here no bonding issues were observed over a period of 10 months.

### 3.2. EAP activity on Hippo-MEA and pathological structural features

The samples analyzed with Hippo-MEA showed varying EAP activity in response to the 4-AP treatment (Fig. 1F-K). Case 1 had 4/59 active electrodes (electrodes detecting EAPs; Fig. 1 F), case 2 1/59 (Fig. 1G) and case 3 4/59 (Fig. 1H). Mean firing rates in active electrodes during the peak drug effect period (970 s to 1270 s) were 0.046 Hz in case 1, 0.010 Hz in case 2 and 0.067 Hz in case 3. In individual electrodes firing rates could momentarily reach as high as 3.333 Hz (Fig. 1H).

The localization of active electrodes (Fig. 1 I-K) did not correspond to the pyramidal cell layer (Fig. 1 L-N), which concurs with the observed loss of NeuN positive cells (neurons) in the pyramidal cell layer and with the pathologist’s assessment of hippocampal sclerosis (Table 1, Histopathology) (Blümcke et al., 2013). Instead, the position of active electrodes corresponds more to the granular cell layer of the dentate gyrus where NeuN positive neurons were evident (Fig.1I vs. L, J vs. M and K vs. N). In addition to sclerosis, hippocampal gliosis is a typical pathological feature of TLE (Blümcke et al., 2013). Extensive gliosis was evident from staining against the astrocytic protein GFAP in all three hippocampi (Fig. 1L-N), which likely contributes to large areas of the slices displaying no EAPs (Fig. 1I-K) (Verkhratsky et al., 2023). It is also possible that the activity levels are affected by the long recovery period (see Sample transport and preparation). Overall, results show that Hippo-MEA can identify active and inactive areas and detect EAPs from sclerotic human hippocampal slices.

### 3.3. LFPs on Hippo-MEA

Similar to the EAP activity, the LFP activity on Hippo-MEA was variable between patients. Representative LFP traces are shown in Fig. 2A and B (patient 3, EAP raster plots included). The frequency band with the highest number of responding electrodes was the delta band (9/59 on average), while the lowest number of responding electrodes was at the alpha band (2/59 on average) (Fig. 2C). The number of electrodes responding on the gamma band was variable (from 2/59 to 11/59). Delta band power increase in electroencephalogram data is a marker of epileptic activity in patients with TLE (De Stefano et al., 2022). Therefore, it is interesting that 4-AP, a known convulsant (Heuzeroth et al., 2019) increased especially delta band power, even in case 2 which was otherwise less active than the others.

**Fig. 2.**
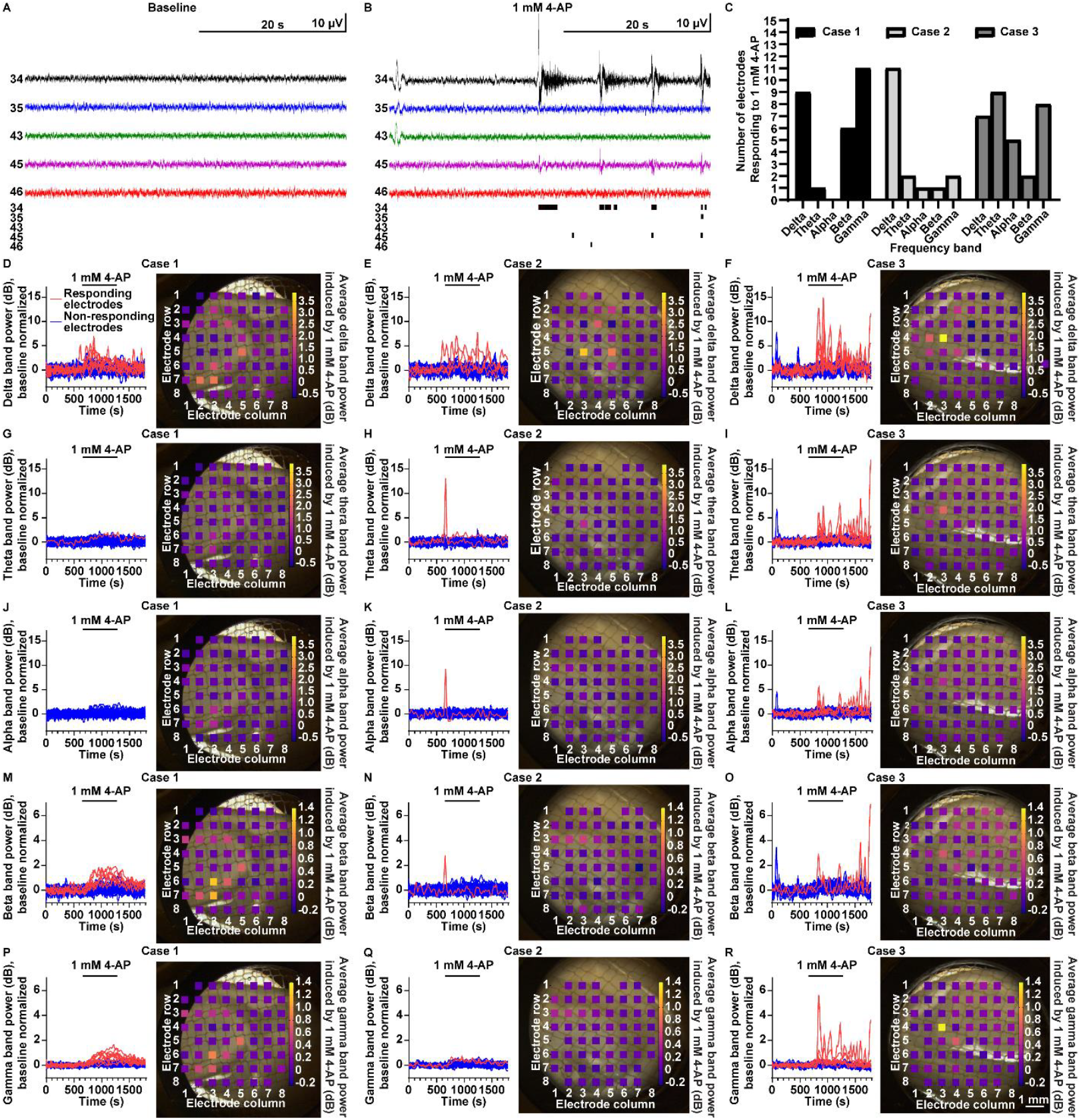
Increased LFP activity on Hippo-MEA in response to 1 mM 4-AP. **A and B**, Representative 200 Hz low-pass filtered LFP traces (upper rows) and corresponding raster plots of spiking activity (lower rows) from 5 electrodes during baseline (A) and 4-AP treatment (B). **C**, Numbers of electrodes responding to 4-AP treatment at each frequency band and each of the three cases. **D, E and F**, 4-AP induced increase in delta band power in cases 1, 2 and 3, respectively. **G, H and I**, 4-AP induced increase in delta band power. J, K and L, 4-AP induced increase in alpha band power. **M, N and O**, 4-AP induced increase in beta band power. **P, Q and R**, 4-AP induced increase in gamma band power. The band power in each electrode was normalized to baseline. In D-R the left panel shows the LFP band power in each electrode as a function of time in 10 s bins, and the right panel shows the average increase in LFP band power during 4-AP treatment across the hippocampal slice. In the case 2 electrodes 51 and 87 were omitted due to noise. Please note the different scales for delta, theta, and alpha bands (D-L) vs. beta and gamma bands (M-R).

The normalized LFP band powers over time are shown in Fig. 2D-R (left), and the average band power increases during 4-AP treatment across the slice in the same subfigures (right). Again, the electrodes responding to the 4-AP stimulation the most were not localized near the pyramidal cell layer, a feature which is likely tied to the sclerosis of pyramidal neurons (Fig. 2D-R vs. Fig. 1L-N; Table 1). Instead, the electrodes showing 4-AP induced increases in LFP activity were closer to the granular cell layer of the dentate gyrus. The results show that the Hippo-MEA detects LFP activity in sclerotic human hippocampal slices and distinguishes the active and inactive sites in the sample.

## 4. Conclusions

The Hippo-MEA and the Sample Cup have suitable dimensions for recording large samples such as human hippocampal slices. The Hippo-MEA can record both EAP and LFP data from different subfields of the human hippocampus simultaneously. While the recording system allows only 59 recording electrodes, limiting the spatial resolution of the array, the recording system is commercially available and already widely used, making Hippo-MEA easily adoptable. Through comparison to data from other methods, the Hippo-MEA can provide a starting point for improving the identification of the optimal resection area in TLE surgeries and understanding the contribution of hippocampal pathology to TLE.

## Author contributions: CRediT

AP, Conceptualization, Data curation, Formal analysis, Investigation, Writing – original draft.

VLez, Investigation, Project administration, Writing – original draft.

TRy, Conceptualization, Formal analysis, Investigation, Resources, Writing – review and editing.

JKr, Investigation, Resources, Writing – review and editing.

VS, Investigation, Writing – review and editing.

MGB, Project administration, Writing – review and editing.

ADP, Project administration, Writing – review and editing.

AD, Methodology, Resources, Writing – review and editing.

JKy, Project administration, Writing – review and editing.

MTA, Data curation, Project administration, Writing – review and editing.

HE, Data curation, Project administration, Writing – review and editing.

TRa, Data curation, Formal analysis, Writing – review and editing.

AI, Methodology, Project administration, Writing – review and editing.

AS, Funding acquisition, Project administration, Resources, Supervision, Writing – review and editing. VLei, Project administration, Resources, Supervision, Writing – review and editing.

RK, Funding acquisition, Project administration, Resources, Supervision, Writing – review and editing. TM, Conceptualization, Funding acquisition, Project administration, Resources, Supervision, Writing – review and editing.

## Data availability

The data that support the findings of this study are not openly available due to reasons of sensitivity. Pseudonymized data may be available from University of Eastern Finland and Kuopio University Hospital, Wellbeing Services County of North Savo for qualified academic investigators after signing a material transfer agreement. Data will be shared for the sole purpose of replicating procedures and results.

## Acknowledgments

The authors are very thankful to the patients who participated in the study and the people who contributed to our Patient and Public Involvement activities. We also thank the whole multidisciplinary Kuopio Epilepsy Center Epilepsy Surgery Group (Kuopio University Hospital) and all multiscale epilepsy project members at the University of Eastern Finland (especially the sample transport team). Thank you to Mirka Tikkanen for cryosectioning the IHC samples. The electrophysiological measurements were enabled by the In vitro and ex vivo electrophysiology core facility (University of Eastern Finland, Biocenter Kuopio and Biocenter Finland). Computational resources were provided by the UEF Bioinformatics Center (University of Eastern Finland).

